# Mixed Model with Correction for Case-Control Ascertainment Increases Association Power

**DOI:** 10.1101/008755

**Authors:** Tristan Hayeck, Noah A. Zaitlen, Po-Ru Loh, Bjarni Vilhjalmsson, Samuela Pollack, Alexander Gusev, Jian Yang, Guo-Bo Chen, Michael E. Goddard, Peter M. Visscher, Nick Patterson, Alkes L. Price

**Affiliations:** Department of Biostatistics, Harvard School of Public Health, Boston, MA; Program in Medical and Population Genetics, Broad Institute of Harvard and MIT, Cambridge, MA; Department of Medicine, Lung Biology Center, University of California, San Francisco, CA; Department of Epidemiology, Harvard School of Public Health, Boston, MA; Queensland Brain Institute, University of Queensland, Brisbane, Australia; University of Queensland Diamantina Institute, University of Queensland, Brisbane, Australia; Faculty of Land and Food Resources, University of Melbourne, Melbourne, Australia

**Author notes:** Correspondence should be addressed to T.H. or A.L.P.

## Abstract

We introduce a Liability Threshold Mixed Linear Model (LTMLM) association statistic for ascertained case-control studies that increases power vs. existing mixed model methods, with a well-controlled false-positive rate. Recent work has shown that existing mixed model methods suffer a loss in power under case-control ascertainment, but no solution has been proposed. Here, we solve this problem using a chi-square score statistic computed from posterior mean liabilities (PML) under the liability threshold model. Each individual’s PML is conditional not only on that individual’s case-control status, but also on every individual’s case-control status and on the genetic relationship matrix obtained from the data. The PML are estimated using a multivariate Gibbs sampler, with the liability-scale phenotypic covariance matrix based on the genetic relationship matrix (GRM) and a heritability parameter estimated via Haseman-Elston regression on case-control phenotypes followed by transformation to liability scale. In simulations of unrelated individuals, the LTMLM statistic was correctly calibrated and achieved higher power than existing mixed model methods in all scenarios tested, with the magnitude of the improvement depending on sample size and severity of case-control ascertainment. In a WTCCC2 multiple sclerosis data set with >10,000 samples, LTMLM was correctly calibrated and attained a 4.1% improvement (P = 0.007) in chi-square statistics (vs. existing mixed model methods) at 75 known associated SNPs, consistent with simulations. Larger increases in power are expected at larger sample sizes. In conclusion, an increase in power over existing mixed model methods is available for ascertained case-control studies of diseases with low prevalence.

## Introduction

Mixed model association statistics are a widely used approach to correct for population structure and cryptic relatedness in genome-wide association studies (GWAS)^1–11^. However, recent work shows that existing mixed model association statistics suffer a loss in power relative to standard logistic regression in ascertained case-control studies^11^. It is widely known that appropriate modeling of case-control ascertainment can produce substantial increases in power for case-control studies with fixed-effect covariates^12–14^, but such increases in power have not yet been achieved with models that include random effects.

We developed an association score statistic based on a liability threshold mixed linear model (LTMLM). The LTMLM statistic relies on the posterior mean liability (PML) of each individual; the PML is calculated using a multivariate Gibbs sampler^15^. The PML of each individual is conditional on the genetic relationship matrix (GRM), the case-control status of every individual, and the disease prevalence. Existing methods use a univariate prospective model to compute association statistics, but here we use a multivariate retrospective model.

The LTMLM statistic provides an increase in power in simulations based on either simulated or real genotypes. In a WTCCC2 multiple sclerosis data set with >10,000 samples, LTMLM was correctly calibrated and attains a 4.1% improvement (P = 0.007) in chi-square statistics (vs. existing mixed model methods) at 75 known associated SNPs, consistent with simulations.

## Materials and Methods

### Overview of Method

We improve upon standard mixed model methods^11^ using a retrospective association score statistic (LTMLM) computed from posterior mean liabilities (PML) under the liability threshold model. The improvement over previous approaches comes from appropriate modeling of case-control ascertainment. We consider all individuals simultaneously, incorporating prevalence information.

Our method consists of three steps. First, the genetic relationship matrix (GRM) is calculated and a corresponding heritability parameter is estimated, modeling the phenotype covariance of all individuals (see *Estimation of Heritability Parameter*). The heritability parameter is estimated using Haseman-Elston (H-E) regression on the observed scale followed by transformation to liability scale. Second, Posterior Mean Liabilities (PML) are estimated using a truncated multivariate normal Gibbs sampler (see *Posterior Mean Liabilities*). The PML of each individual is conditional on that individual’s case-control status, on every other individual’s case-control status, and on disease prevalence and liability-scale phenotypic covariance. Third, a chi-square (1 d.o.f) association score statistic is computed based on the association between the candidate SNP and the PML (see *LTMLM Association Statistic*).

The toy example in Figure 1 provides an illustration of how genetic relatedness to a disease case can increase an individual’s PML. In Figure 1a and 1b, we plot the distribution of liabilities in 10,000 unrelated individuals with random ascertainment and case-control ascertainment (for a disease with prevalence 0.1%), respectively. In Figure 1c and 1d, we plot the same distributions conditional on an individual having genetic relatedness of 0.5 to a disease case, assuming liability-scale heritability of 1.0. In each case, the posterior distribution of liabilities (and hence the PML) is shifted upwards. (The magnitude and direction of this effect would be different for an individual having a genetic relatedness of 0.5 to a control.) Our main focus below is on much lower levels of genetic relatedness (identity-by-state) among many unrelated samples, but the same principles apply.

**Figure 1.**
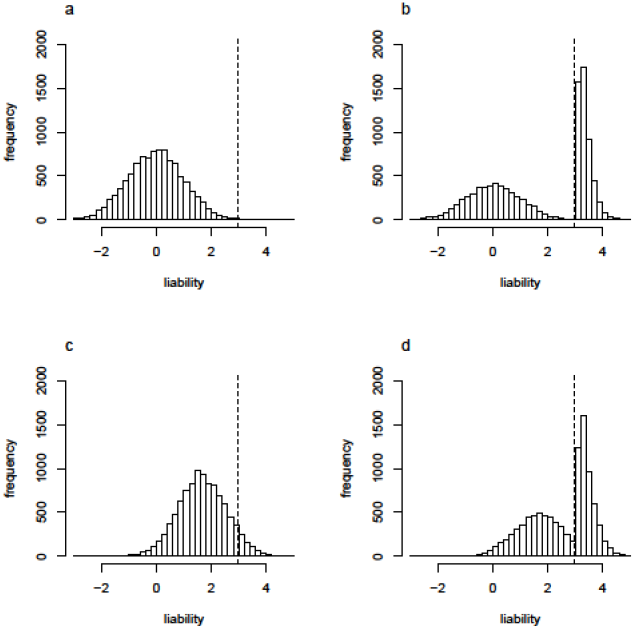
Genetic relatedness to a disease case can increase an individual’s PML. In (a) and (b), we plot distributions of liabilities for a set of 10,000 individuals under (a) random ascertainment or (b) case-control ascertainment for a disease with prevalence 0.1% (see Figure 2 of Lee et al.^17^). In (c) and (d), we plot the same distributions conditional on an individual having genetic relatedness of 0.5 to a disease case, assuming a heritability of 1 on the liability scale.

### Estimation of Heritability Parameter

Mixed model association statistics rely on the estimation of a heritability parameter. We note that this heritability parameter, which Kang et al.^4^ referred to as “pseudo-heritability”, is generally lower than the total narrow-sense heritability (*h*^2^) in data sets not dominated by family relatedness, but may be larger than the heritability explained by genotyped SNPs (*h_g_*^2^)^16^ in data sets with population structure or family relatedness. However, for ease of notation, we use the symbol *h*^2^ to represent this heritability parameter. A list of all notation used below is provided in Table S1.

The goal is to test for association between a candidate SNP and a phenotype. We first consider a quantitative trait:

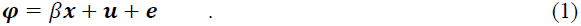

The phenotypic data (transformed to have mean 0 and variance 1) may be represented as a vector ***φ*** with values for each individual *i*. Genotype values of candidate SNP are transformed to a vector ***x*** with mean 0 and variance 1, with effect size *β*. The quantitative trait value depends on the fixed effect of the candidate SNP (*β**x***), the genetic random effect excluding the candidate SNP (***u*),** and the environmental component (***e***). We extend to case-control traits via the liability threshold model, in which each individual has an underlying, unobserved normally distributed trait called the liability. An individual is a disease case if the liability exceeds a specified threshold *t*, corresponding to disease prevalence^17^ (Figure S1).

Standard mixed model association methods generally estimate *h*^2^ from a genetic relationship matrix (GRM) and phenotypes using restricted maximum likelihood (ReML) ^4; 11^. Genotypic data is used to build a GRM (excluding the candidate SNP^11^):

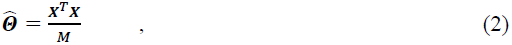

where ***X*** is a matrix of non-candidate SNPs normalized to mean 0 and variance 1 and *M* is the number of SNPs. We estimate *h*^2^ using Haseman-Elston (H-E) regression followed by a transformation to liability scale. The H-E regression estimate is obtained by regressing the product of the case-control phenotypes on the off diagonal terms of the GRM^18–20^:

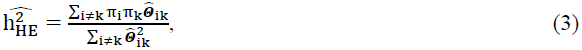

where *π_i_* denotes the case-control status of individual *i* and 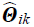 is the genetic relatedness of individuals *i* and *k*. This gives an estimate on the observed scale which is then transformed to the liability scale^21^:

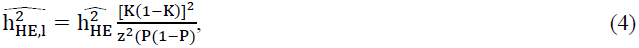

where *z* is the height of the standard normal density 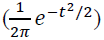 at the liability threshold *t*, *K* is disease prevalence, and *P* is the proportion of cases in the sample^21^.

Then, the variance between the individuals is modeled as the phenotypic covariance

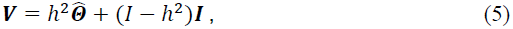

Where 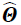 the *N* by *N* GRM, ***V*** is the phenotypic covariance, *h*^2^ is the heritability parameter, and ***I*** is the identity matrix.

Using the phenotypic covariance matrix ***V***, the liability is modeled as a multivariate normal distribution:

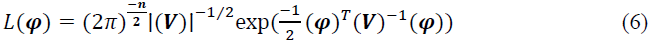

We note that we observe the case-control phenotypes of the individuals and not the continuous liabilities.

### Posterior Mean Liabilities

We first consider the univariate PML (PMLuni), constructed independently for each individual; we generalize to the multivariate setting below. As described in equations 11 and 12 of ref. ^21^, these correspond to the expected value of the liability conditional on the case control status:

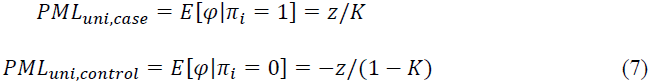

These values are calculated analytically in the univariate setting, and can be thought of as the mean of a truncated normal above or below the liability threshold *t* depending on case control status^21^.

We now consider the multivariate PML (PML_multi_), estimated jointly across individuals. The PML_multi_ for each individual is conditional on that individual’s case-control status, on every other individual’s case-control status, and on their phenotypic covariance. The PML_multi_ is estimated using a Gibbs sampler, analogous to previous work^15^ (which focused on family relatedness and did not consider association statistics). The Gibbs sampler is an iterative algorithm that generates random variables from conditional distributions in order to avoid the difficult task of explicitly calculating the marginal density for each random variable.

For each individual in turn, the conditional distribution of the liability is calculated based on all of the other individuals and a new value is generated. The algorithm is:

Initialization: for each individual *j*,

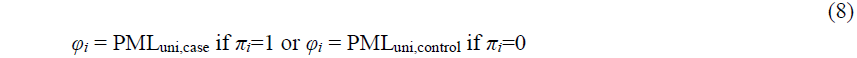

For each MCMC iteration *n*

For each individual *i*

Sample *φ_i_* from the constrained conditional univariate normal distribution L(*φ_i_*) ∼ exp(–*φ*^T^V ^-1^*φ*/2) and constraint *φ_i_*≥*t* if *π_i_*=1, *φ_i_*<*t* if *π_i_*=0 (where *φ*_≠_*_i_* are fixed)

We use 100 burn-in iterations followed by 1,000 additional MCMC iterations. We estimate the PML*_multi_* by averaging over MCMC iterations. We reduce the number of MCMC iterations needed via Rao-Blackwellization, which averages (across iterations *n*) the *posterior means* of the distributions from which each *φ_i_* is sampled.

### LTMLM Association Statistic

The LTMLM association statistic is calculated using PML_multi_. For simplicity, we first consider the case where the liability is known. We jointly model the liability and the genotypes using a retrospective model, enabling appropriate treatment of sample ascertainment. We concatenate the two vectors (***φ***,***x***) and derive the joint likelihood for these combined terms. The covariance of ***φ*** and ***x*** between individual i and k is:

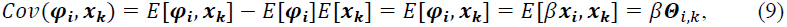

where Θ is the true underlying genetic relatedness matrix from which genotypes are sampled. (We note that Θ, which is unobserved, is different from the GRM 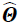 estimated from the data.) The variance of (***φ***,***x***) as a function of effect size *β* is:

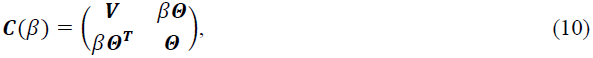

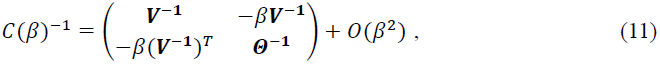

where both of these matrices are 2N by 2N. The joint likelihood of the liability and genotypes are distributed as a multivariate normal *N*(0,***C***(*β*)), and thus

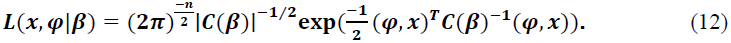

Taking the derivative of the log likelihood results in the score equation. The determinant of the matrix ***V*** does not have any terms linear in ***β***, so the terms with ***V*** alone drop out when we take the derivative:

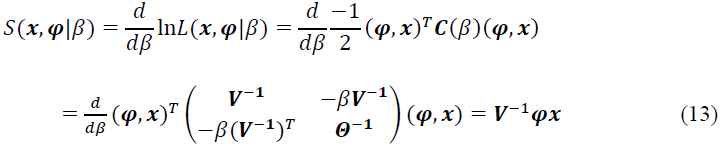

The marginal score statistic tests the null hypothesis that the fixed effect of the candidate SNP is zero (H0: *β* = 0) vs. the alternative hypothesis (HA: *β* ≠ 0). The denominator of the score statistic is the variance of the score evaluated under the null. :

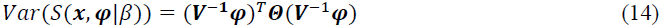

This leads to the score statistic:

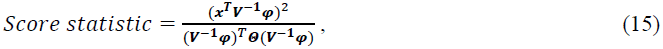

where ***Θ***, the true underlying genetic relatedness of the individuals, can be approximated by the identity matrix in data sets of unrelated individuals.

We now consider a case-control trait, with unobserved liability, and derive the score function using the observed case-control status of each individual, ***π***. Returning to the score function and conditioning on case control status:

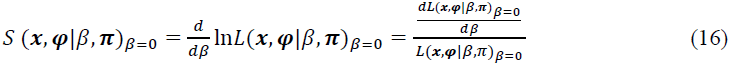

Introducing the unobserved quantitative liability, *φ*, the score function can be rewritten in terms of the probability density of the liability:

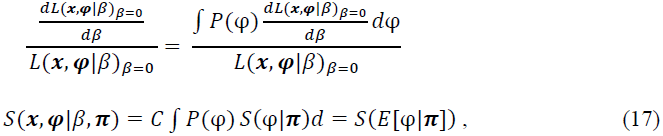

where *P*(*φ*) is the probability density of the liability and *E*[*φ| π*] is the PML. It follows that an appropriate score statistic is

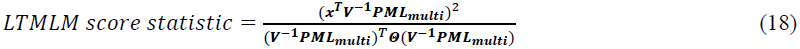

Again ***Θ*** can be approximated by the identity matrix in data sets of unrelated individuals; we note that this choice affects only a constant calibration factor (since the denominator is the same for each candidate SNP), and that other calibration options are available (see below). As with other association statistics, the LTMLM score statistic generalizes to non-normally distributed genotypes^22–24^. The overall computational cost of computing the LTMLM statistic is O(*MN^2^*) when *M* > *N* (Table S2).

We calculate the GRM via Leave One Chromosome Out (LOCO) analysis, i.e. for each candidate SNP on a given chromosome the GRM is calculated using all of the other chromosomes. This prevents deflation due to double counting the candidate SNP as both a fixed effect and random effect in the mixed model^4; 6; 11^.

### Simulated Genotypes and Simulated Phenotypes

We performed simulations both using simulated genotypes and simulated phenotypes, and using real genotypes and simulated phenotypes (see below). Quantitative liabilities for each individual were generated from SNP effects and an environmental component. The proportion of causal SNPs was set to 0.01. The quantitative liabilities were then dichotomized based on the liability threshold to categorize each individual as a case or control. Case-control ascertainment was performed, simulating 50% cases and 50% controls. We compared ATT, MLM, and LTMLM statistics (see Table 1). MLM statistics were computed using the GCTA-LOCO statistic described in ref. ^11^, with the heritability parameters estimated using the GCTA software^25^. All simulations used *M* SNPs to calculate the GRM and *M* additional SNP as candidate SNPs being tested for association (so that candidate SNPs are not included in the GRM).

**Table 1.** List of association statistics. We list properties of the Armitage Trend Test (ATT), standard mixed model association statistic (MLM), and proposed statistic (LTMLM). π* is normalized case-control status (mean 0, variance 1), x are normalized genotypes, PML_uni_ is the univariate PML conditional on the case-control status of a single individual, PML_multi_ is the multivariate PML conditional of the case-control status of all individuals, I is the identity matrix, V is the phenotypic covariance (on the observed scale for MLM, and on the liability scale for LTMLM).

|  | ATT | MLM | LTMLM |
| --- | --- | --- | --- |
| **Quant vs Case control Trait** | Both | Both | CC |
| **Herit Param Est.** | None | ReML | H-E regression |
| **Prospective vs. Retrospective** | Pro | Pro | Retro |
| **Equation** | $\frac{(x^T \pi^*)^2}{x^T x}$ | $\frac{(x^T \pi^* V^{-1})^2}{x^T V^{-1} x} = \frac{(x^T V^{-1} PML_{uni})^2}{x^T V^{-1} x}$ | $\frac{(x^T V^{-1} PML_{multi})^2}{(V^{-1} PML_{multi})^T I (V^{-1} PML_{multi})}$ |
| **Corrects for Confounding?** | No | Yes | Yes |
| **Models Case-Control Asc.** | No | No | Yes |

In the primary analyses, we simulated individuals without population structure or LD, with *N* = 1 K or 5 K samples, *M* = 1 K, 5 K or 50 K SNPs, and prevalence *K* = 50%, 10%, 1% or 0.1%. Genotypes were sampled from independent binomials with allele frequencies uniform on [0.1,0.9]. In secondary analyses, we simulated population structure by simulating two populations with an *F*_ST_ of 0.01, whose allele frequencies were drawn from beta distributions with parameters *p*(1 – *F*_ST_)/ *F*_ST_ and (1 – *p*)(1 – *F*_ST_)/ *F*_ST_, based on ancestral allele frequency *p* which is uniform on [0.1,0.9].

### WTCCC2 Genotypes and Simulated Phenotypes

We also conducted simulations using real genotypes from WTCCC2 to incorporate LD and realistic population structure. The WTCCC2 data contained 360,557 SNPs and 15,633 samples, as described previously^11^. Since the goal of the power study is demonstrate a comparison of the statistics under case-control ascertainment, we used *N* = 1000 samples (500 cases and 500 controls), with simulated phenotypes having prevalence of 50%, 25%, 10%. The prevalence was restricted to a lower bound of 10% because of the limitation of only 15,633 WTCCC2 samples for simulating case-control ascertainment. We computed ATT, MLM and LTMLM statistics as described above.

### WTCCC2 Genotypes and MS Phenotypes

Finally, we analyzed WTCCC2 individuals with ascertained case-control phenotypes for MS^11^, a disease with a prevalence of around 0.1%. We computed ATT, MLM and LTMLM statistics as described above. Although the underlying MS study was appropriately matched for ancestry^26^, the data made available to researchers included only pan-European cases and UK controls. Thus, the WTCCC2 data set shows a severe mismatch in ancestry of cases and controls; this severe mismatch between cases and controls is not representative of a typical GWAS. We thus restricted our primary analysis to 10,034 samples with only a moderate mismatch in ancestry, but analyses of unmatched and stringently matched data sets were also performed (Figure S2). The unmatched data set contained 10,204 case and 5,429 controls. Matching was performed by first calculating 20 PCs in the full cohort and weighing the contribution of each PC based on the variance in phenotype it explained in a multiple regression. A Euclidean distance over these 20 weighted dimensions was then computed for all pairs of individuals, and each case was greedily assigned the nearest unmatched control until no matched case-control pairs could be identified. Finally, any matched case-control pairs that were not within 6 standard deviations of the mean pairwise distance were removed as outliers, yielding the 5,017 cases and 5,017 matched controls used in our primary analysis. Stringent matching was performed by additionally removing any matched case-control pairs that were not within 2 standard deviations of the mean pairwise distance, yielding 4,094 cases and 4,094 matched controls used in our stringently matched analysis.

We compared association statistics at 75 published SNPs associated to MS^11^. We used a jackknife approach to assess the statistical significance of differences in association statistics, by excluding each of the 75 published SNPs in turn.

## Results

### Simulations: Simulated Genotypes and Simulated Phenotypes

We first conducted simulations using simulated genotypes and simulated ascertained case-control phenotypes (see Materials and Methods). Our main simulations involve unrelated individuals with no population structure, but the impact of population structure is explored below. We evaluated the power of ATT, MLM and LTMLM (see Table 2). Results for additional values of #SNPs (*M*) and #samples (*N*) are displayed in Table S3. The LTMLM statistic consistently outperforms the ATT and MLM statistics, particularly at low values of disease prevalence. For LTMLM vs. MLM at disease prevalence of 0.1%, 3% and 24% improvements were observed in simulations with 5,000 SNPs and 50,000 SNPs respectively. Smaller improvements were observed at higher disease prevalences. Test statistics were well-calibrated at null markers. Simulations at other values of *M* and *N* indicate that the magnitude of the improvement depends on the value of *N*/*M* (Table S3). Simulations with population structure demonstrate similar results, but with inflation in the ATT statistic as expected (Table S4).

**Table 2.** Results on simulated genotypes and simulated phenotypes. We report average χ^2^ statistics. *N* is the number of individuals and *M* is the number of SNPs. SNP set indicates either all SNPs, the 1% causal SNPs, or the 99% null SNPs. The disease prevalence ranges from 50% (no case-control ascertainment) to 0.1%.

| M | N | PREVALENCE | SNP SET | ATT | MLM | LTMLM |
| --- | --- | --- | --- | --- | --- | --- |
| 5000 | 5000 | 50% | all | 1.14 | 1.15 | 1.15 |
|  |  |  | causal | 16.79 | 17.13 | 17.12 |
|  |  |  | null | 0.99 | 0.99 | 0.99 |
|  |  | 10% | all | 1.23 | 1.23 | 1.24 |
|  |  |  | causal | 24.59 | 24.79 | 25.18 |
|  |  |  | null | 0.99 | 0.99 | 0.99 |
|  |  | 1% | all | 1.45 | 1.40 | 1.46 |
|  |  |  | causal | 45.72 | 42.61 | 46.85 |
|  |  |  | null | 1.00 | 0.99 | 1.00 |
|  |  | 0.1% | all | 1.71 | 1.51 | 1.73 |
|  |  |  | causal | 71.99 | 59.53 | 74.05 |
|  |  |  | null | 1.00 | 0.93 | 1.00 |
| 50000 | 5000 | 50% | all | 1.02 | 1.02 | 1.02 |
|  |  |  | causal | 2.68 | 2.68 | 2.69 |
|  |  |  | null | 1.00 | 1.00 | 1.00 |
|  |  | 10% | all | 1.02 | 1.02 | 1.02 |
|  |  |  | causal | 3.35 | 3.35 | 3.35 |
|  |  |  | null | 1.00 | 1.00 | 1.00 |
|  |  | 1% | all | 1.04 | 1.04 | 1.04 |
|  |  |  | causal | 5.47 | 5.42 | 5.48 |
|  |  |  | null | 1.00 | 1.00 | 1.00 |
|  |  | 0.1% | all | 1.07 | 1.07 | 1.07 |
|  |  |  | causal | 8.02 | 7.81 | 8.05 |
|  |  |  | null | 1.00 | 1.00 | 1.00 |

The MLM statistics were calculated using an *h*^2^ parameter estimated using Restricted Maximum Likelihood Methods (ReML)^4^, but the LTMLM statistics were calculated using an *h^2^* parameter estimated via Haseman-Elston (H-E) regression on case-control phenotypes followed by transformation to liability scale ^18; 21^ (see Materials and Methods). As case-control ascertainment becomes more severe the H-E regression estimate of the *h^2^* remains unbiased, whereas the variance component estimate is severely downwardly biased even after transformation to the liability scale (Table 3 and Table S5), consistent with previous work (see ref.^19^ and Supp Table 9 of ref.^11^) . Population structure resulted in bias of both ReML and HE-regression estimates of *h^2^*, but consistently higher bias for the ReML estimates (Table S6). These biases do not inflate LTMLM or MLM statistics under the null (Table S4). We note that previous work has shown that running MLM using the correct *h^2^* parameter does not ameliorate the loss in power for MLM^11^.

**Table 3.**
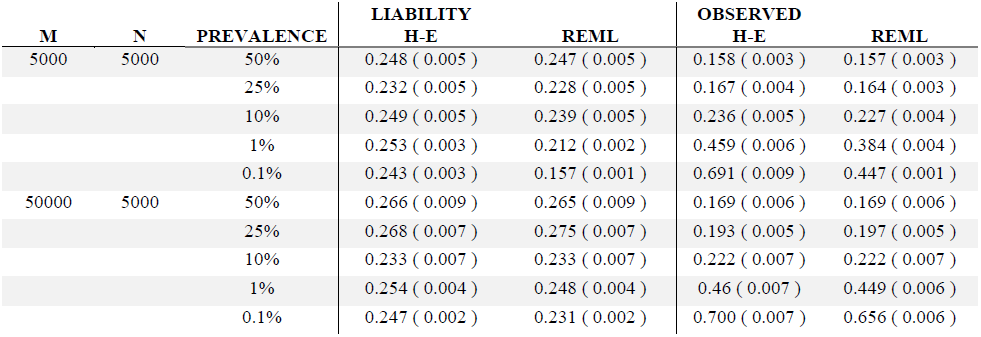
Heritability parameter estimates on simulated genotypes and phenotypes. These results are from the same simulations used to generate Table 2. We report results on both liability and observed scales. The true *h^2^* explained by the SNPs used to build the GRM is 25% on the liability scale for all simulations.

### Simulations: WTCCC2 Genotypes and Simulated Phenotypes

We next conducted simulations using real WTCCC2 genotypes and simulated ascertained case-control phenotypes (see Materials and Methods)^11; 26^. For a given value of *M* (*M* SNPs to calculate the GRM and *M* candidate SNPs, for a total of 2*M* SNPs), we used the first *M*/2 SNPs from each of the first four chromosomes. The GRM was calculated using SNPs on chromosomes 3 and 4, with SNPs on chromosomes 1 and 2 treated as the candidate SNPs. The simulated phenotypes were generated from chromosome 1 and 3, where 1% of the SNPs were randomly selected as being causal. Results are reported for causal SNPs on chromosome 1 and null SNPs on chromosome 2, which were not used to build the GRM.

Results for 1,000 and 10,000 SNPs (*M*) are displayed in Table 4 and Table S7, with sample size fixed at 500 cases and 500 controls; formal power calculations produce similar results (Table S8). Once again, the LTMLM statistic outperforms ATT and MLM as case-control ascertainment becomes more severe. (A limitation of these simulations is that performing case-control ascertainment on a fixed set of individuals limits case-control sample size; thus, these simulations were restricted to a disease prevalence of 10% or higher. It is reasonable to infer that for rarer diseases with more extreme case-control ascertainment the LTMLM statistic would achieve even higher power gains, as was demonstrated in simulations with simulated genotypes.)

**Table 4.** Results on real genotypes and simulated phenotypes. We report average χ^2^ statistics. *M* is the number of SNPs, and sample size is fixed at 500 cases and 500 controls.

| M | PREVALENCE | SNP SET | ATT | MLM | LTMLM |
| --- | --- | --- | --- | --- | --- |
| 1000 | 50% | all | 1.61 | 1.64 | 1.64 |
|  |  | causal | 17.07 | 17.96 | 17.90 |
|  |  | null | 1.01 | 1.01 | 1.01 |
|  | 25% | all | 1.73 | 1.75 | 1.75 |
|  |  | causal | 19.64 | 20.46 | 20.56 |
|  |  | null | 1.02 | 1.01 | 1.01 |
|  | 10% | all | 1.90 | 1.89 | 1.93 |
|  |  | causal | 24.88 | 25.15 | 26.11 |
|  |  | null | 1.04 | 1.02 | 1.03 |
| 10000 | 50% | all | 1.08 | 1.08 | 1.08 |
|  |  | causal | 2.65 | 2.66 | 2.66 |
|  |  | null | 1.00 | 1.00 | 1.00 |
|  | 25% | all | 1.10 | 1.10 | 1.10 |
|  |  | causal | 2.90 | 2.91 | 2.92 |
|  |  | null | 1.02 | 1.01 | 1.01 |
|  | 10% | all | 1.14 | 1.13 | 1.13 |
|  |  | causal | 3.58 | 3.58 | 3.61 |
|  |  | null | 1.03 | 1.02 | 1.02 |

The *h*^2^ parameter estimates for simulations using real genotypes are displayed in Table S9. The H-E regression estimates are unbiased, but the ReML estimates are again downwardly biased at lower prevalence and large *N*/*M*.

### WTCCC2 Multiple Sclerosis data set

We analyzed the WTCCC2 genotypes together with multiple sclerosis (MS) case-control phenotypes: 5,172 MS cases and 5,172 controls genotyped on Illumina chips^11; 26^ (see Materials and Methods). We compared ATT, ATT with 5 PCs (PCA)23, MLM and LTMLM. We evaluated calibration using the average χ^2^ over all SNPs; we note that the average χ^2^ over all SNPs is expected to be greater than 1 due to polygenic effects^11; 27^, and all methods can be correctly calibrated via LD Score regression^28^.

We evaluated power using the average χ^2^ over the 75 published SNPs. The results are displayed in Table 5. The LTMLM method performed best, with a 4.1% improvement vs. MLM (jackknife *P* = 0.007; see Materials and Methods) and an even larger improvement versus ATT and PCA, consistent with simulations (Table 2). Similar results are obtained when calibrating association statistics via LD Score regression^28^ (Table S10). A perfectly matched data set with 4,094 MS cases and 4,094 controls yielded a similar improvement for LTMLM vs. MLM (Table S11). We also applied LTMLM to the full unmatched data set of 10,204 MS cases and 5,429 controls, where there is a severe mismatch in ancestry between cases and controls that is not representative of a typical GWAS. The LOCO estimates of *h^2^* demonstrate inflation before controlling for population structure (Table S12). In this analysis, the H-E regression estimate of the *h^2^* produces an unrealistic value of 7.3 on the observed scale (corresponding to 2.8 on the liability scale), which is outside the plausible 0-1 range suggesting severe population stratification or other severe problems with the data. We do not recommend the use of LTMLM on unmatched samples when such severe problems are detected. For completeness, we report the results of running LTMLM, which results in a loss in power (Table S11).

**Table 5.** **Results on WTCCC2 MS data set.** We report the genome**-**wide average χ^2^ over 360,557 SNPs and the average across 75 published SNPs, before or after normalizing by the genome-wide average. All results are based on analysis of 10,034 individuals (see main text).

| SNP SET | ATT | PCA | MLM | LTMLM |
| --- | --- | --- | --- | --- |
| **Genome Wide Average** | 1.38 | 1.16 | 1.14 | 1.17 |
| **Published SNPs Average** | 11.64 | 9.97 | 9.92 | 10.59 |
| **Published SNPs/Genome Wide Average** | 8.44 | 8.61 | 8.67 | 9.03 |

## Discussion

We have shown that controlling for case-control ascertainment using the LTMLM statistic can lead to significant power improvements in ascertained case-control studies of diseases of low prevalence. This was demonstrated via simulations using both simulated and real genotypes, and in WTCCC2 MS case-control data.

The LTMLM statistic should not be used if the inferred liability-scale *h^2^* parameter is outside the plausible 0-1 bound, as this is indicative of severe population stratification or other severe problems with the data (this can also be assessed via PCA; see Figure S2). In such settings, either matching based on ancestry should first be performed, or other statistics should be used.

Several limitations of LTMLM remain as directions for future study. First, previous work has shown that using the posterior mean liabilities in conjunction with fixed effects such as BMI, age, or known associated SNPs will further increase power^12; 21^. The incorporation of fixed-effect covariates into the LTMLM statistic is not considered here, and remains as a future direction. Second, the calibration of our statistic in unrelated samples relies on an approximation that works well in the WTCCC2 data analyzed, but may not work well in all data sets. Here, calibration via LD Score regression offers an appealing alternative^28^. Third, we did not consider ascertained case-control studies in family data sets, which also represents a future direction. Fourth, LTMLM requires running time O(*MN*^2^), analogous to standard mixed model association methods. This may be computationally intractable in very large data sets. We are developing much faster mixed model methods^29^, but those methods do not consider case-control ascertainment and should not be applied to ascertained case-control data for diseases of low prevalence. The incorporation of the ideas we have described here into those methods is an open question. Finally, our methods could potentially be extended to multiple traits^7; 30; 31^.

## Acknowledgements

We thank D. Golan and S. Rosset for helpful discussions. This research was funded by NIH grant R01 HG006399.

This study makes use of data generated by the Wellcome Trust Case Control Consortium (WTCCC). A full list of the investigators who contributed to the generation of the data is available from http://www.wtccc.org.uk/.

## Web Resources

The URLs for data presented herein are as follows:

1. Liability Threshold Mixed Linear Model (LTMLM) software will be provided at http://www.hsph.harvard.edu/alkes-price/software/
2. GCTA (Genome-wide Complex Trait Analysis) software, http://www.complextraitgenomics.com/software/gcta/

---

**Figure S1.**
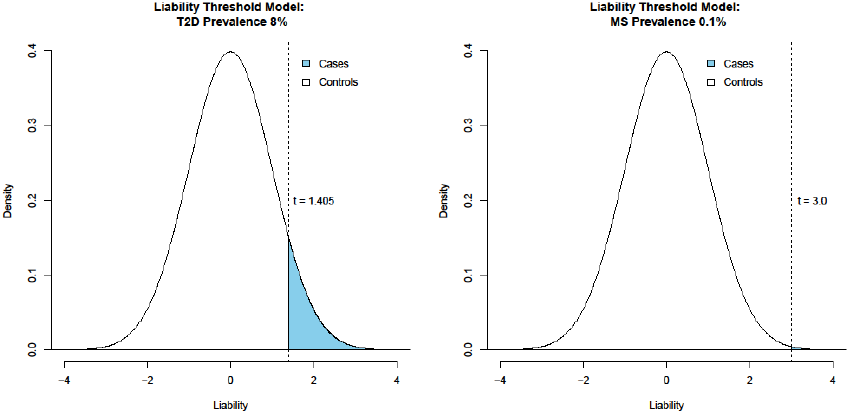
Liability Threshold Model. The liability threshold model performs a transformation based on disease prevalence. As ascertainment becomes more drastic so does the difference between the PML for cases versus controls. In Figure S1, the portion of the population above the threshold is a case (blue). For T2D, at a prevalence of 8% (blue), the threshold is set to 1.405. In this region, the expected value for the posterior liability is 1.85 and the expected value for the controls is −0.14. Comparing T2D to MS with disease prevalence around 0.1% and t around 3.00, the PML_indiv_ for a control is 0.00 and 3.33 for a case. As the disease prevalence goes down the difference in the PML_indiv_ for cases versus controls increases, the transformation plays a larger role for rare diseases and results in a power gain for the LTMLM.

**Figure S2.**
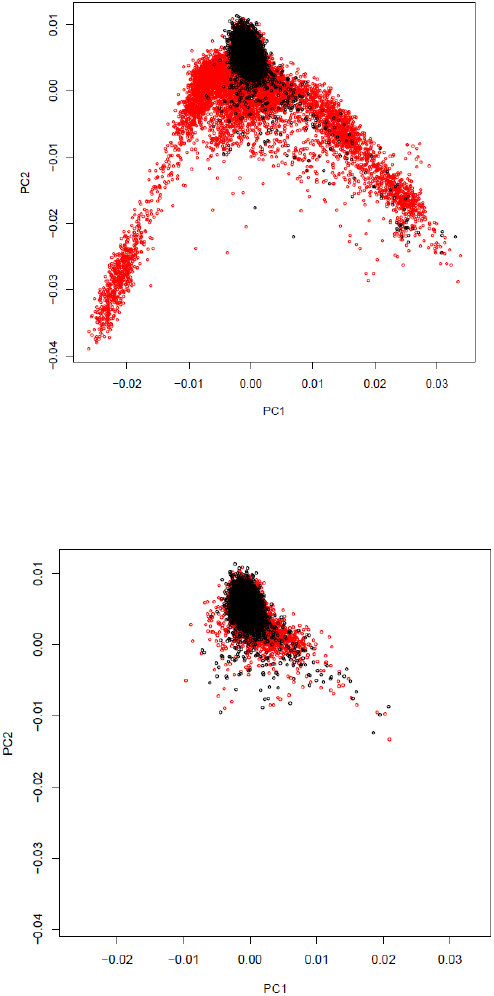
Mismatch in ancestry between MS cases and controls. We plot the first two principal components for (a) unmatched data with a severe mismatch (5,429 MS cases and 10,204controls), (b) stringently matched data using the first 20 PC(4,094 MS cases and 4,094 controls). The controls are depicted in red and cases in black. After PC matching the remaining samples show considerably less population stratification differentiation between cases and controls.

**Table S1:** Description of notation used and a brief description of the terms.

| **Term** | **Description** |
| --- | --- |
| $\phi$ | Quantitative liability, the unobserved trait |
| $\beta$ | Effect Size of the SNP |
| $x$ | Genotype values of candidate SNP, normalized to mean 0 variance 1 |
| $u$ | Genetic random effect excluding the candidate SNP |
| $e$ | Environmental component |
| $X$ | Matrix of genotype values of non-candidate SNPs, normalized to mean 0 and variance 1 |
| $\pi$ | Observed binary case control phenotype. |
| $t$ | Threshold corresponding to the disease prevalence |
| $K$ | Prevalence of the disease in the population |
| $P$ | Proportion of cases in the sample |
| $\hat{\Theta}$ | Genetic Relationship Matrix (GRM) computed from the data |
| $\Theta$ | True underlying Genetic Relationship Matrix (GRM) |
| $V$ | Phenotypic covariance matrix |
| $I$ | Identity matrix |
| $h^2$ | Heritability parameter |

**Table S2.** Computational cost. *M* is the number of SNPs and *N* is the number of individuals. We assume that *M* >*N*. The details of the computational costs of MLM are provided in Table 1 of ref^11^.

| **Computation** | **ATT** | **MLM** | **LTMLM** |
| --- | --- | --- | --- |
| **GRM and $V^{-1}$** | NA | $O(MN^2)$ | $O(MN^2)$ |
| **PML** | NA | NA | $O(N^3 + iter * N^2) = O(N^3)$ |
| **Assoc. Statistic** | $O(MN)$ | $O(MN)$ or $O(MN^2)$ | $O(MN)$ |
| **Overall** | $O(MN)$ | $O(MN^2)$ | $O(MN^2)$ |

**Table S3.**
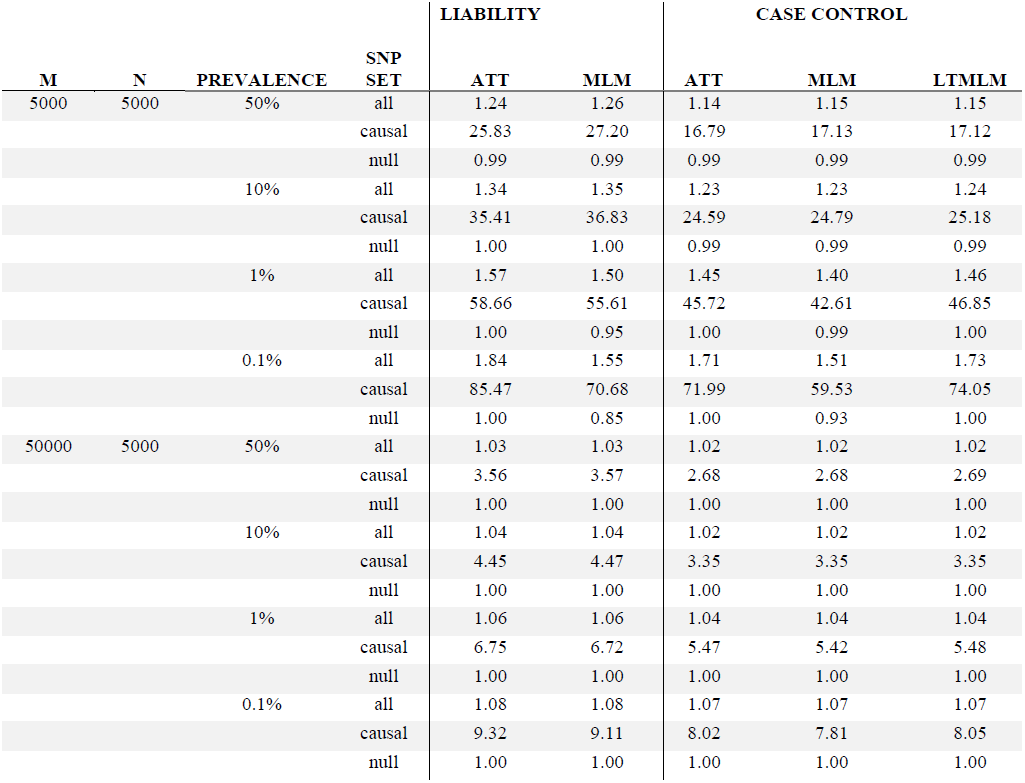
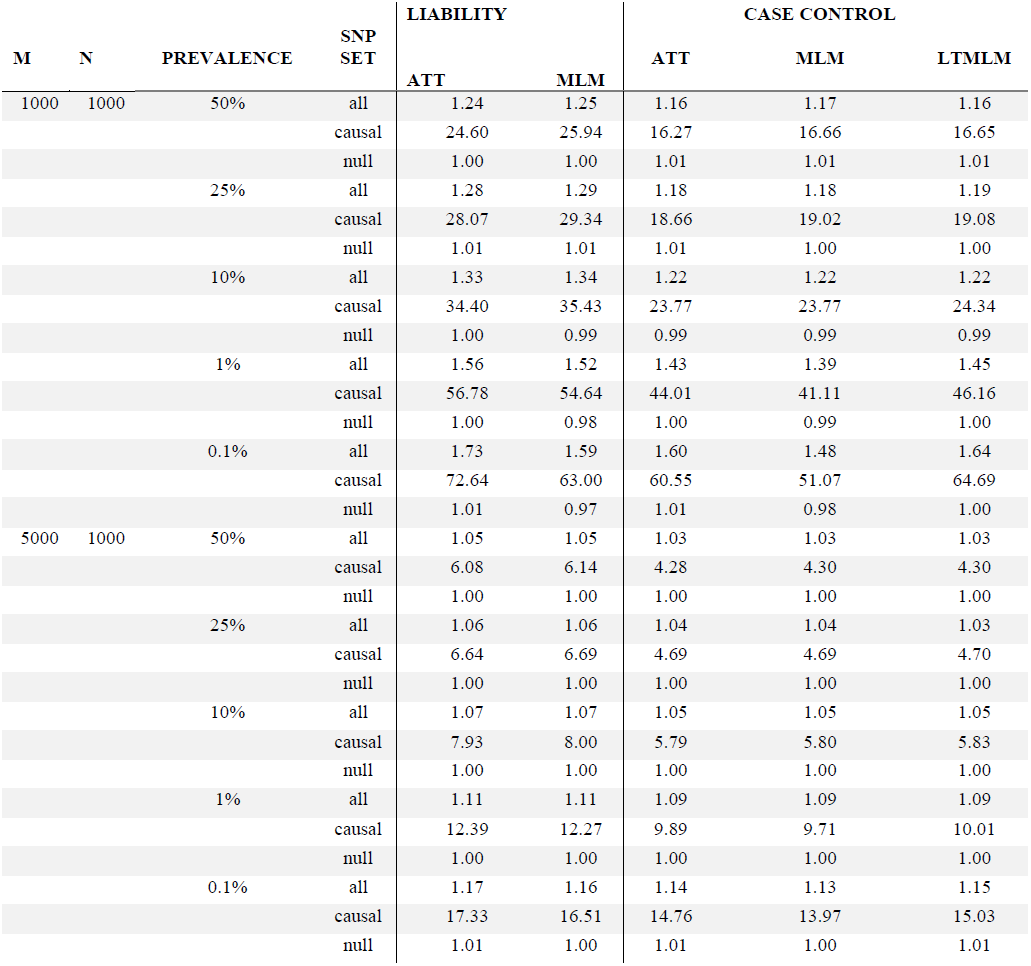
Complete results on simulated genotypes and simulated phenotypes. Results are analogous to Table 2, but are reported for other values of *M* and *N*. For completeness, we also report ATT and MLM statistics computed using the underlying liability, where we again observe a loss in power for MLM at lower prevalence.

| M | N | PREVALENCE | SNP<br>SET | LIABILITY |  | CASE CONTROL |  |  |
| --- | --- | --- | --- | --- | --- | --- | --- | --- |
|  |  |  |  | ATT | MLM | ATT | MLM | LTMLM |
| 5000 | 5000 | 50% | all | 1.24 | 1.26 | 1.14 | 1.15 | 1.15 |
|  |  |  | causal | 25.83 | 27.20 | 16.79 | 17.13 | 17.12 |
|  |  |  | null | 0.99 | 0.99 | 0.99 | 0.99 | 0.99 |
|  |  | 10% | all | 1.34 | 1.35 | 1.23 | 1.23 | 1.24 |
|  |  |  | causal | 35.41 | 36.83 | 24.59 | 24.79 | 25.18 |
|  |  |  | null | 1.00 | 1.00 | 0.99 | 0.99 | 0.99 |
|  |  | 1% | all | 1.57 | 1.50 | 1.45 | 1.40 | 1.46 |
|  |  |  | causal | 58.66 | 55.61 | 45.72 | 42.61 | 46.85 |
|  |  |  | null | 1.00 | 0.95 | 1.00 | 0.99 | 1.00 |
|  |  | 0.1% | all | 1.84 | 1.55 | 1.71 | 1.51 | 1.73 |
|  |  |  | causal | 85.47 | 70.68 | 71.99 | 59.53 | 74.05 |
|  |  |  | null | 1.00 | 0.85 | 1.00 | 0.93 | 1.00 |
| 50000 | 5000 | 50% | all | 1.03 | 1.03 | 1.02 | 1.02 | 1.02 |
|  |  |  | causal | 3.56 | 3.57 | 2.68 | 2.68 | 2.69 |
|  |  |  | null | 1.00 | 1.00 | 1.00 | 1.00 | 1.00 |
|  |  | 10% | all | 1.04 | 1.04 | 1.02 | 1.02 | 1.02 |
|  |  |  | causal | 4.45 | 4.47 | 3.35 | 3.35 | 3.35 |
|  |  |  | null | 1.00 | 1.00 | 1.00 | 1.00 | 1.00 |
|  |  | 1% | all | 1.06 | 1.06 | 1.04 | 1.04 | 1.04 |
|  |  |  | causal | 6.75 | 6.72 | 5.47 | 5.42 | 5.48 |
|  |  |  | null | 1.00 | 1.00 | 1.00 | 1.00 | 1.00 |
|  |  | 0.1% | all | 1.08 | 1.08 | 1.07 | 1.07 | 1.07 |
|  |  |  | causal | 9.32 | 9.11 | 8.02 | 7.81 | 8.05 |
|  |  |  | null | 1.00 | 1.00 | 1.00 | 1.00 | 1.00 |

**Table S3. Complete results on simulated genotypes and simulated phenotypes.**
| M | N | PREVALENCE | SNP SET | LIABILITY |  | CASE CONTROL |  |  |
| --- | --- | --- | --- | --- | --- | --- | --- | --- |
|  |  |  |  | ATT | MLM | ATT | MLM | LTMLM |
| 1000 | 1000 | 50% | all | 1.24 | 1.25 | 1.16 | 1.17 | 1.16 |
|  |  |  | causal | 24.60 | 25.94 | 16.27 | 16.66 | 16.65 |
|  |  |  | null | 1.00 | 1.00 | 1.01 | 1.01 | 1.01 |
|  |  | 25% | all | 1.28 | 1.29 | 1.18 | 1.18 | 1.19 |
|  |  |  | causal | 28.07 | 29.34 | 18.66 | 19.02 | 19.08 |
|  |  |  | null | 1.01 | 1.01 | 1.01 | 1.00 | 1.00 |
|  |  | 10% | all | 1.33 | 1.34 | 1.22 | 1.22 | 1.22 |
|  |  |  | causal | 34.40 | 35.43 | 23.77 | 23.77 | 24.34 |
|  |  |  | null | 1.00 | 0.99 | 0.99 | 0.99 | 0.99 |
|  |  | 1% | all | 1.56 | 1.52 | 1.43 | 1.39 | 1.45 |
|  |  |  | causal | 56.78 | 54.64 | 44.01 | 41.11 | 46.16 |
|  |  |  | null | 1.00 | 0.98 | 1.00 | 0.99 | 1.00 |
|  |  | 0.1% | all | 1.73 | 1.59 | 1.60 | 1.48 | 1.64 |
|  |  |  | causal | 72.64 | 63.00 | 60.55 | 51.07 | 64.69 |
|  |  |  | null | 1.01 | 0.97 | 1.01 | 0.98 | 1.00 |
| 5000 | 1000 | 50% | all | 1.05 | 1.05 | 1.03 | 1.03 | 1.03 |
|  |  |  | causal | 6.08 | 6.14 | 4.28 | 4.30 | 4.30 |
|  |  |  | null | 1.00 | 1.00 | 1.00 | 1.00 | 1.00 |
|  |  | 25% | all | 1.06 | 1.06 | 1.04 | 1.04 | 1.03 |
|  |  |  | causal | 6.64 | 6.69 | 4.69 | 4.69 | 4.70 |
|  |  |  | null | 1.00 | 1.00 | 1.00 | 1.00 | 1.00 |
|  |  | 10% | all | 1.07 | 1.07 | 1.05 | 1.05 | 1.05 |
|  |  |  | causal | 7.93 | 8.00 | 5.79 | 5.80 | 5.83 |
|  |  |  | null | 1.00 | 1.00 | 1.00 | 1.00 | 1.00 |
|  |  | 1% | all | 1.11 | 1.11 | 1.09 | 1.09 | 1.09 |
|  |  |  | causal | 12.39 | 12.27 | 9.89 | 9.71 | 10.01 |
|  |  |  | null | 1.00 | 1.00 | 1.00 | 1.00 | 1.00 |
|  |  | 0.1% | all | 1.17 | 1.16 | 1.14 | 1.13 | 1.15 |
|  |  |  | causal | 17.33 | 16.51 | 14.76 | 13.97 | 15.03 |
|  |  |  | null | 1.01 | 1.00 | 1.01 | 1.00 | 1.01 |

**Table S4.** Results on simulated genotypes and simulated phenotypes with population structure. We report average χ^2^ statistics for simulations with population structure (see main text).

| M | N | PREVALENCE | SNP SET | ATT | MLM | LTMLM |
| --- | --- | --- | --- | --- | --- | --- |
| 1000 | 1000 | 10% | All | 1.99 | 1.32 | 1.25 |
|  |  |  | Causal | 24.28 | 16.59 | 16.03 |
|  |  |  | Null | 1.54 | 1.01 | 0.95 |
|  |  |  | Causal/All | 12.19 | 12.54 | 12.80 |
| 10000 |  |  | All | 1.13 | 1.04 | 1.03 |
|  |  |  | Causal | 3.40 | 3.24 | 3.20 |
|  |  |  | Null | 1.08 | 1.00 | 0.99 |
|  |  |  | Causal/All | 3.02 | 3.11 | 3.10 |
| 20000 |  |  | All | 1.10 | 1.02 | 1.01 |
|  |  |  | Causal | 2.26 | 2.16 | 2.13 |
|  |  |  | Null | 1.07 | 1.00 | 0.99 |
|  |  |  | Causal/All | 2.07 | 2.11 | 2.10 |

**Table S5.**
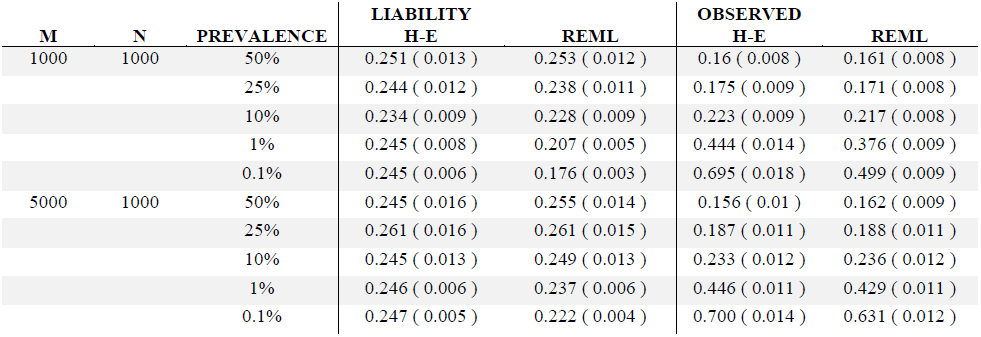
Heritability parameter estimates on simulated genotypes and phenotypes. Results are analogous to Table 3, under different settings of *M* and *N*.

| M | N | PREVALENCE | LIABILITY |  | OBSERVED |  |
| --- | --- | --- | --- | --- | --- | --- |
|  |  |  | H-E | REML | H-E | REML |
| 1000 | 1000 | 50% | 0.251 ( 0.013 ) | 0.253 ( 0.012 ) | 0.16 ( 0.008 ) | 0.161 ( 0.008 ) |
|  |  | 25% | 0.244 ( 0.012 ) | 0.238 ( 0.011 ) | 0.175 ( 0.009 ) | 0.171 ( 0.008 ) |
|  |  | 10% | 0.234 ( 0.009 ) | 0.228 ( 0.009 ) | 0.223 ( 0.009 ) | 0.217 ( 0.008 ) |
|  |  | 1% | 0.245 ( 0.008 ) | 0.207 ( 0.005 ) | 0.444 ( 0.014 ) | 0.376 ( 0.009 ) |
|  |  | 0.1% | 0.245 ( 0.006 ) | 0.176 ( 0.003 ) | 0.695 ( 0.018 ) | 0.499 ( 0.009 ) |
| 5000 | 1000 | 50% | 0.245 ( 0.016 ) | 0.255 ( 0.014 ) | 0.156 ( 0.01 ) | 0.162 ( 0.009 ) |
|  |  | 25% | 0.261 ( 0.016 ) | 0.261 ( 0.015 ) | 0.187 ( 0.011 ) | 0.188 ( 0.011 ) |
|  |  | 10% | 0.245 ( 0.013 ) | 0.249 ( 0.013 ) | 0.233 ( 0.012 ) | 0.236 ( 0.012 ) |
|  |  | 1% | 0.246 ( 0.006 ) | 0.237 ( 0.006 ) | 0.446 ( 0.011 ) | 0.429 ( 0.011 ) |
|  |  | 0.1% | 0.247 ( 0.005 ) | 0.222 ( 0.004 ) | 0.700 ( 0.014 ) | 0.631 ( 0.012 ) |

**Table S6.**
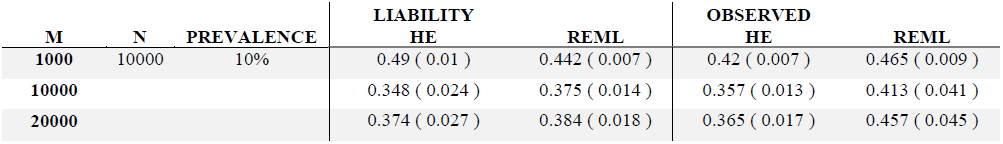
Heritability parameter estimates on simulated genotypes and phenotypes with population structure. These results are from the same simulations used to generate Table S5. We report results on both liability and observed scales. The true *h^2^* explained by the SNPs used to build the GRM is 25% on the liability scale for all simulations. We report results on both liability and observed scales. The true *h^2^* explained by the SNPs used to build the GRM is 25% on the liability scale for all simulations.

| M | N | PREVALENCE | LIABILITY |  | OBSERVED |  |
| --- | --- | --- | --- | --- | --- | --- |
|  |  |  | HE | REML | HE | REML |
| **1000** | 10000 | 10% | 0.49 ( 0.01 ) | 0.442 ( 0.007 ) | 0.42 ( 0.007 ) | 0.465 ( 0.009 ) |
| **10000** |  |  | 0.348 ( 0.024 ) | 0.375 ( 0.014 ) | 0.357 ( 0.013 ) | 0.413 ( 0.041 ) |
| **20000** |  |  | 0.374 ( 0.027 ) | 0.384 ( 0.018 ) | 0.365 ( 0.017 ) | 0.457 ( 0.045 ) |

**Table S7.**
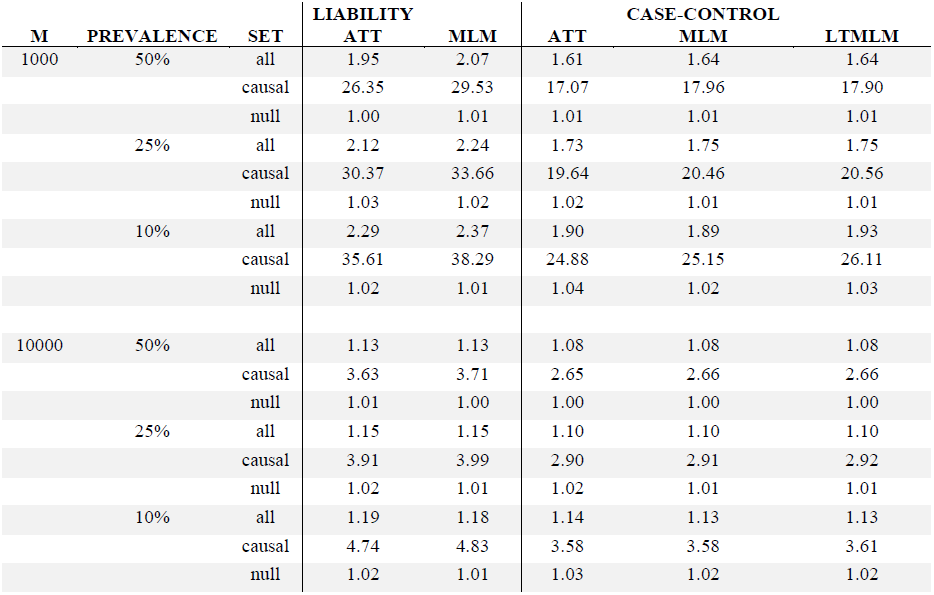
Complete results on real genotypes and simulated phenotypes. Results are analogous to Table 4, but we also report ATT and MLM statistics computed using the underlying liability.

**Table S8.** Percentage of causal SNPs achieving genome-wide significance (p < 5 × 10^−8^). Results are based on 500 cases and 500 controls with real genotypes and simulated phenotypes, where *M* = 1000 SNPs.

| PREVALENCE | ATT | MLM | LTMLM |
| --- | --- | --- | --- |
| 50% | 30.0% | 29.7% | 30.4% |
| 25% | 21.4% | 22.2% | 22.5% |
| 10% | 21.4% | 22.2% | 22.5 |

**Table S9.**
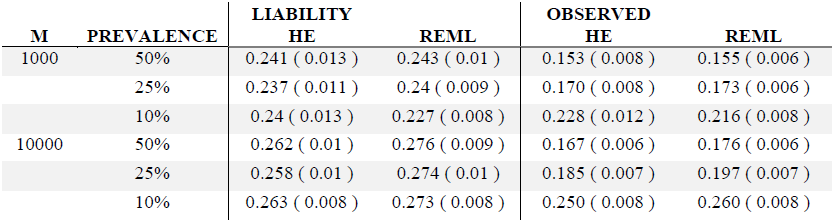
Heritability parameter estimates on real genotypes and simulated phenotypes. These results are from the same simulations used to generate Table 4. The true heritability explained by the SNPs used to build the GRM is 25% on the liability scale for all simulations.

| M | PREVALENCE | LIABILITY |  | OBSERVED |  |
| --- | --- | --- | --- | --- | --- |
|  |  | HE | REML | HE | REML |
| 1000 | 50% | 0.241 ( 0.013 ) | 0.243 ( 0.01 ) | 0.153 ( 0.008 ) | 0.155 ( 0.006 ) |
|  | 25% | 0.237 ( 0.011 ) | 0.24 ( 0.009 ) | 0.170 ( 0.008 ) | 0.173 ( 0.006 ) |
|  | 10% | 0.24 ( 0.013 ) | 0.227 ( 0.008 ) | 0.228 ( 0.012 ) | 0.216 ( 0.008 ) |
| 10000 | 50% | 0.262 ( 0.01 ) | 0.276 ( 0.009 ) | 0.167 ( 0.006 ) | 0.176 ( 0.006 ) |
|  | 25% | 0.258 ( 0.01 ) | 0.274 ( 0.01 ) | 0.185 ( 0.007 ) | 0.197 ( 0.007 ) |
|  | 10% | 0.263 ( 0.008 ) | 0.273 ( 0.008 ) | 0.250 ( 0.008 ) | 0.260 ( 0.008 ) |

**Table S10:** Results on WTCCC2 MS data set with calibration via LD Score regression. We report the genome wide χ^2^ averages using 10,034 individuals over 360,557 SNPs and the average across 75 published SNPs standardized by the genome wide average and LD Score regression intercept.

| SNP SET | ATT | PCA | MLM | LTMLM |
| --- | --- | --- | --- | --- |
| GENOME WIDE AVERAGE | 1.38 | 1.16 | 1.14 | 1.17 |
| GENOME WIDE LD SCORE INTERCEPT | 1.29 | 1.09 | 1.08 | 1.10 |
| PUBLISHED SNPS AVERAGE | 11.64 | 9.97 | 9.92 | 10.59 |
| PUBLISHED SNPS/GENOME WIDE AVERAGE | 8.44 | 8.61 | 8.67 | 9.03 |
| PUBLISHED SNPS/LD SCORE INTERCEPT | 9.06 | 9.17 | 9.20 | 9.66 |

**Table S11.** Results on WTCCC2 MS data set at different levels of QC. We report results for stringently matched (*N* = 8,188), partially matched (*N* = 10,034) and unmatched (*N* = 15,633) data sets (see main text).

| **SNP SET** | **N** | **ATT** | **MLM** | **LTMLM** |
| --- | --- | --- | --- | --- |
| GENOME WIDE AVERAGE | 8188 | 1.16 | 1.11 | 1.14 |
| PUBLISHED SNPS AVERAGE |  | 8.94 | 8.26 | 8.76 |
| PUBLISHED SNPS/GENOME WIDE AVERAGE |  | 7.73 | 7.45 | 7.71 |
| GENOME WIDE AVERAGE | 10034 | 1.38 | 1.14 | 1.17 |
| PUBLISHED SNPS AVERAGE |  | 11.64 | 9.92 | 10.59 |
| PUBLISHED SNPS/GENOME WIDE AVERAGE |  | 8.44 | 8.67 | 9.03 |
| GENOME WIDE AVERAGE | 15633 | 3.95 | 1.23 | 1.08 |
| PUBLISHED SNPS AVERAGE |  | 18.54 | 11.30 | 5.76 |
| PUBLISHED SNPS/GENOME WIDE AVERAGE |  | 4.69 | 9.20 | 5.32 |

**Table S12:** Heritability parameter estimates on WTCCC2 MS data set at different levels of QC. We report results for stringently matched (*N* = 8,188), partially matched (*N* = 10,034) and unmatched (*N* = 15,633) data sets (see main text).

| **N** | **Liability** |  | **Observed** |  |
| --- | --- | --- | --- | --- |
|  | **HE** | **ReML** | **HE** | **ReML** |
| **8188** | 0.363 (0.0017) | 0.260 (0.001) | 0.979 (0.005) | 0.702 (0.003) |
| **10034** | 0.704 (0.009) | 0.279 (0.001) | 1.901 (0.025) | 0.753 (0.002) |
| **15633** | 2.792 (0.010) | 0.293 (0.001) | 7.543 (0.0266) | 0.792 (0.002) |

